# Application of Global Metabolomics to the Identification of Complex Counterfeit Medicinal Products

**DOI:** 10.1101/567339

**Authors:** Conor Jenkins, Ben Orsburn

**Affiliations:** Think20 Labs, Columbia, MD; Hood College Department of Biology, Frederick, MD 21702; Proteomics und Genomics, Baltimore, MD

## Abstract

Food fraud and drug counterfeiting are of increasingly large concern to both global economics and to public health and safety. Simple medicinal products consisting of single synthesized or purified compounds can be tested for purity and authenticity rapidly with established assays such as chromatography and UV absorbance. Drugs derived from natural sources may contain hundreds or thousands of distinct chemical compounds and require correspondingly complex analytical methods. In this study we explore the use of methods developed for global metabolic profiling toward the identification of unknown complex medicinal products. By utilizing rapid solvent extraction followed by ultrahigh pressure high performance liquid chromatography (UHPLC) coupled to high resolution accurate mass spectrometry (HRAM-MS/MS), we can reliably obtain a profile of the sample’s molecular makeup. After profiling plant material to the depth of over 1,000 distinct molecules identified and quantified, we utilize these profiles to identify separately prepared and individually assayed blinded samples. We conclude that once a comprehensive library of small molecules has been acquired for each sample, identical preparations of products of unknown origin may be identified using simple statistical tools such as principal component analysis. We also conclude that these tools will be a valuable resource in affordably identified contaminated, adulterated and counterfeit products.

**Abstract Graphic:** 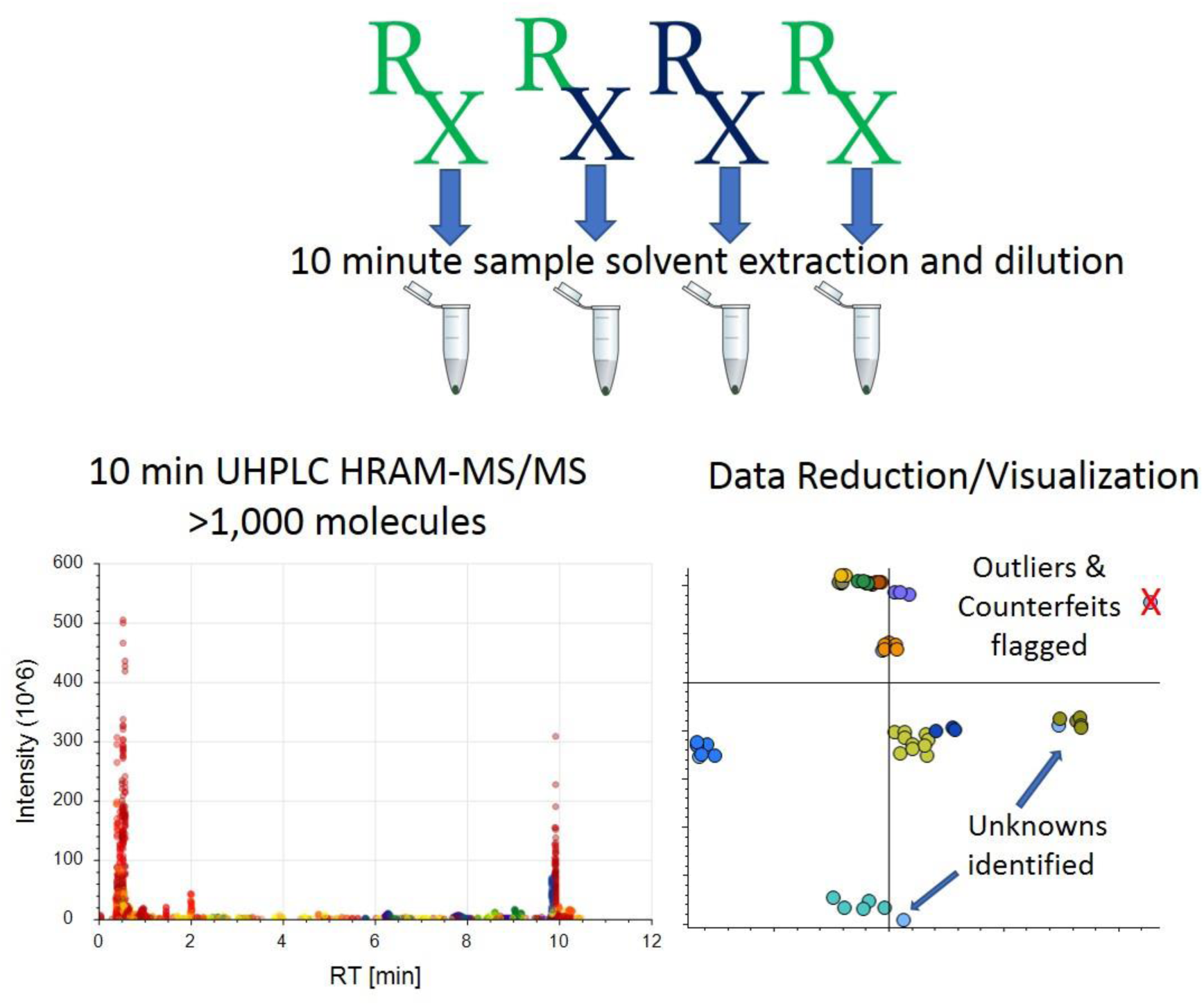

## Introduction

Advances in chromatography and mass spectrometry have led to a massive increase in the number of molecules that can be identified and quantified by these techniques.^1,2^ Recent studies utilizing quadrupole Orbitrap mass spectrometers and multiple chromatography methods have profiled thousands of compounds within single studies. The careful curation of libraries of mass spectra and tandem mass spectra (MS/MS) by groups such as NIST^3^, the Scripps institute through METLIN^4^ and mzCloud, as well as many others has allowed the near instant identification of tens of thousands of compounds by matching experimental MS1 and MS/MS to these curated libraries.

Food fraud is an issue of increasing concern to global economics and for public safety.^5^ A recent analysis estimated that nearly 10% of the global food market is affected in some way by food fraud^6^. A common example of cases appears to consist of the adulteration of legitimate products with materials of lower production cost, such as the use of sweeteners in fruit juices. Adulteration may be a serious concern for people with allergies or other health problems where diet is of paramount concern.^7^

Similarly, drug counterfeiting has been estimated to be in play in up to 10% of the world’s medications, with numbers reaching as high as 30% in parts of the developing world.^8^ The World Health Organization defines a counterfeit drug “as those which have been deliberately and fraudulently mislabeled with respect to identity and/or source. Counterfeit medicines may include medicines with the correct ingredients or with the wrong ingredients, without active ingredients, with insufficient active ingredients or with fake packaging.” Established chemical methods such as high-performance liquid chromatography (HPLC) with ultraviolet (UV) detection can rapidly determine the purity of most simple drugs consisting of one purified or synthesized active product, and is a cost effective solution for detecting mislabeled and adulterated products.^8^ The complexity of natural products such as plant material or compounds that are crudely enriched from biological material is at a level where HPLC and UV detection is simply inadequate, as hundreds or thousands of small molecules may be present in these materials. Furthermore, HPLC may not allow the identification of materials that do possess the correct active materials but were produced or synthesized in an inferior manner by black market chemists. Many new methods have been developed to detect counterfeit alprazolam tablets as black market production can replicate both tablet size, shape and even packaging.^9–11^

With the recent relaxing of many of the North American laws of the 20^th^ century on the plants of the Cannabis genus, many locales are witnessing the introduction of products from these plants. The fibers of the hemp plant are reentering commercial manufacturing and will continue to do so as more North American farms obtain permits for growth and production. Extracts of cannabis plants that lack the intoxicating tetrahydrocannabinol (THC) compounds are being marketed for multiple reasons and ailments. The most obvious change in these laws may be the number of locales that are permitting medical and/or recreational use of Cannabis flowers that do possess high levels of THC. Due to the legal restrictions of the 20^th^ century, little serious work has been performed on these latter materials.^12^

In this study we detail the use of UHPLC-HRAM-MS/MS (uLC-MS/MS) obtain the first comprehensive snapshot of the small molecule features in these plants by retention time, ionization polarity, mass to charge ratio (m/z) and nuclear isotope distribution to a small molecule profile for each of these materials. We search this reference library against multiple openly available MS1 and MS/MS annotated libraries and can assign putative identifications to 390 of the compounds found in these materials. By use of statistical tools developed for metabolomic profiling we find that we can identify separately prepared and blinded samples by matching to those from our known libraries. With a total sample preparation and run time of less than 30 minutes per sample, we conclude that UHPLC HRAM-MS is an attractive solution to the identification of mislabeled, counterfeit and adulterated medicinal products of any level of complexity.

## Materials and Methods

### Samples

Six medical Cannabis samples were obtained by Think20 labs under the guidelines of the MMCC regulations in accordance with a temporary license granted under COMAR 10.62.33.^13^ The strains obtained are listed in Supplemental Table 1 and will be referred to herein as Samples 1 thru 6.

### Sample Preparation

Twenty milligrams of each commercial flower were separated with a surgical steel scalpel in a sterile hood. Compounds were extracted at −20C for 1 hour with 150 µL of buffer consisting of 40% methanol 40% acetonitrile and 20% Milli-Q water. Solid matter was precipitated at 13,000 x g using an Eppendorf 5415C microcentrifuge at 4C for 5 minutes. The top 100 µL of the extract was removed and transferred to a new 1.5 mL tube. In order to clarify the solution and return the solution to baseline LC conditions, the solution was diluted 10x in 4°C 0.1% formic acid in LC-MS grade water. The solution was again centrifuged to precipitate insoluble compounds and the aqueous portion was transferred to glass autosampler vials.

### Blinded Match Samples

Identical samples were prepared from 5 samples as described above, two days following the original extraction. Sample 6 was not prepared as insufficient quantities were available. A second scientist who has requested to remain uncredited removed the samples with scalpel and labeled the tube with an encoded signature. As there was no clear way of discerning visually between the samples, all downstream extraction and processing was performed as described above by the corresponding author.

### UHPLC-MS/MS

The LC-MS system consisted of a Dionex U3000 UHPLC system coupled to a Q Exactive Classic quadrupole Orbitrap system. A simple UHPLC gradient consisting of 0.1% formic acid in Milli-Q water as buffer A and 0.1% formic acid in 100% Acetonitrile as buffer B. A flow rate of 400 µL/minute was used for all experiments. A 10 cm 2.1mm HyperSil Gold 2 µm C-18 column was used for reverse phase separation all experiments. The gradient began at 5% buffer B and ramped to 95% B in 8 minutes, where it maintained for 1 minute before returning to baseline conditions for 2 minutes prior to the next injection. Three technical replicates were performed on day 1 in each polarity followed by a blank consisting of water and the extraction buffer at the same concentration as the experimental samples. The blanks were saved for baseline extraction during data analysis. In all experiments 2 µL of sample was injected directly on column. Source conditions for the mass spectrometer were created automatically within the manufacturer’s Tune software modeled on a 400 µL flow rate. Full mass spectra (MS1) were acquired from 82 – 750 m/z at 70,000 resolution and an ion injection (AGC) target of 5e6 charges per scan. Up to the 5 most intense ions from each MS1 spectra were selected for MS/MS fragmentation if they met the following minimum conditions: >1.6e5 intensity and determined charge stage of 1-3. Ions selected for fragmentation were isolated with a 2.2 Da window symmetrical on the most intense isotope. The ion injection target was 1e5 or 50 milliseconds total ion accumulation time if the target could not be reached. Ions selected for fragmentation were collected as three separate ion packets that were fragmented with a normalized collision energy of 10, 30 and 100, respectively. The three fragmented packets were combined into a single MS/MS spectrum acquired at 17,500 resolution. Automatic gain control was modeled automatically assuming a full width half maximum (FWHM) peak width of 6 seconds. All ions were excluded from a secondary fragmentation event by dynamically excluding all molecules within 10ppm of the fragmented ion for 15 seconds after the event.

### Data Processing

All vendor RAW instrument files were processing in Compound Discoverer 2.1 (CD) from Thermo Fisher. CD processes files through a stepwise logical tree consisting of 9 basic steps. These are detailed fully in Supplemental file #1. Briefly, all files were aligned using a simple linear alignment with a 2 minute mass shift on molecules within a 5ppm mass tolerance. Gaps were imputed according to manufacturer settings and features were searched against the ChemSpider libraries: ACToR: Aggregated Computational Toxicology Resource, FDA UNII – NLM, and FoodDB. Molecular formulas were derived using the in house algorithms and MS/MS spectra were searched against the mZCloud. Feature ideas were assigned preferentially to mzCloud, followed by ChemSpider, and finally by molecular formula.

### Graphical Representation

Figures utilized for data visualization and investigation were generated using R version 3.5.2 (2018-12-20) and are available at https://github.com/jenkinsc11/CD_Metabolomics_Datavis.

## Results and Discussion

### RAW Data Acquisition and Feature ID

The total run time of each single sample injection was 11 minutes. A separate injection was utilized with the mass spectrometer in negative polarity mode being the only difference in the method. Each file was between 100 and 130 MB in size in the vendor proprietary binary format. The samples resulted in the identification of 12,069 chromatographic features in positive ion detection mode and 4,028 chromatographic features in negative mode. An example of the chromatographic features observed in a single RAW file is shown in **Figure 4A**. Useful chromatographic features corresponding to unique Cannabis compounds were obtained across the entire UHPLC gradient. Interestingly, the majority of molecules eluted at high and low points across the gradient and suggest that optimization of the UHPLC may allow further shortening of these experiments with no loss in data quality. Compound Discoverer 2.1 (CD) reduced these to 662 and 404 compounds, respectively. This was performed using the vendor default feature to compound reduction strategies and settings described in Supplemental File 2. As shown in the Venny analysis^14^ in **Figure 5A**, the results of the two polarities were highly complementary, as only 15 named compounds (3.2%) were identified in both polarities. **Figures 5B** and **5C** show the number of compounds that were assigned names using the libraries employed in this study. In both polarities, the majority of compounds were not assigned names, as these masses and/or molecular formulas were not present in these libraries. The mzCloud utilizes a proprietary similarity search algorithm that appears similar to the Hybrid search algorithm recently described by NIST.^15^ By allowing similarity search according to the settings in Supplemental File 2 an additional 144 and 112 compounds were assigned as “similar to” compounds present in the mZCloud library compound when a delta mass shift is allowed, in positive and negative polarities, respectively. In ultra-high-resolution mass spectrometry, molecular formulas may be confidently assigned for relatively small compounds, based purely on mass alone. All but 54 and 96 compounds could be assigned a chemical formula in the negative and positive experiments, respectively. The assignment of these formulas is not fully revealed in this proprietary software, but likely follows the 7 golden rules detailed by Kind and Fiehn.^16^

The accuracy of compound identification depends on the level of match between the library compounds and those of the experimental samples. In an ideal circumstance, the spectral library would contain all possible chemical compounds with MS1, and MS/MS obtained under identical conditions. To the best of our knowledge, no comprehensive small molecule library exists for Cannabis plants. Further analysis using a larger and/or more pertinent set of compound libraries may be necessary to improve the quality of the compound identifications obtained from the files described in this study.

It is also worth noting that the estimation of false discovery rates, has been a historical goal ^17^ and recent focus for small molecule identification.^18,19^ However, these tools are not presently in mainstream use and do not exist in the software used for this study. The mzCloud library does possess a sophisticated scoring mechanism for quality of MS/MS spectra as shown in **Figure 1B** for the amino acid asparagine, but the other libraries rely on MS1 mass accuracy alone. Due to the lack of FDR statistics, several compound identifications appear unlikely to be correct. One example is shown in **Figure 3A**. N-Boc based compound synthesis has never been performed in our newly constructed facility where all sample preparation and analysis has been performed and we find the putative identification an obvious false match to the libraries searched (Data not shown). Manual analysis will be described in conjunction with more sophisticated compound library match tools described in a study by our group elsewhere.

**Figure 1.**
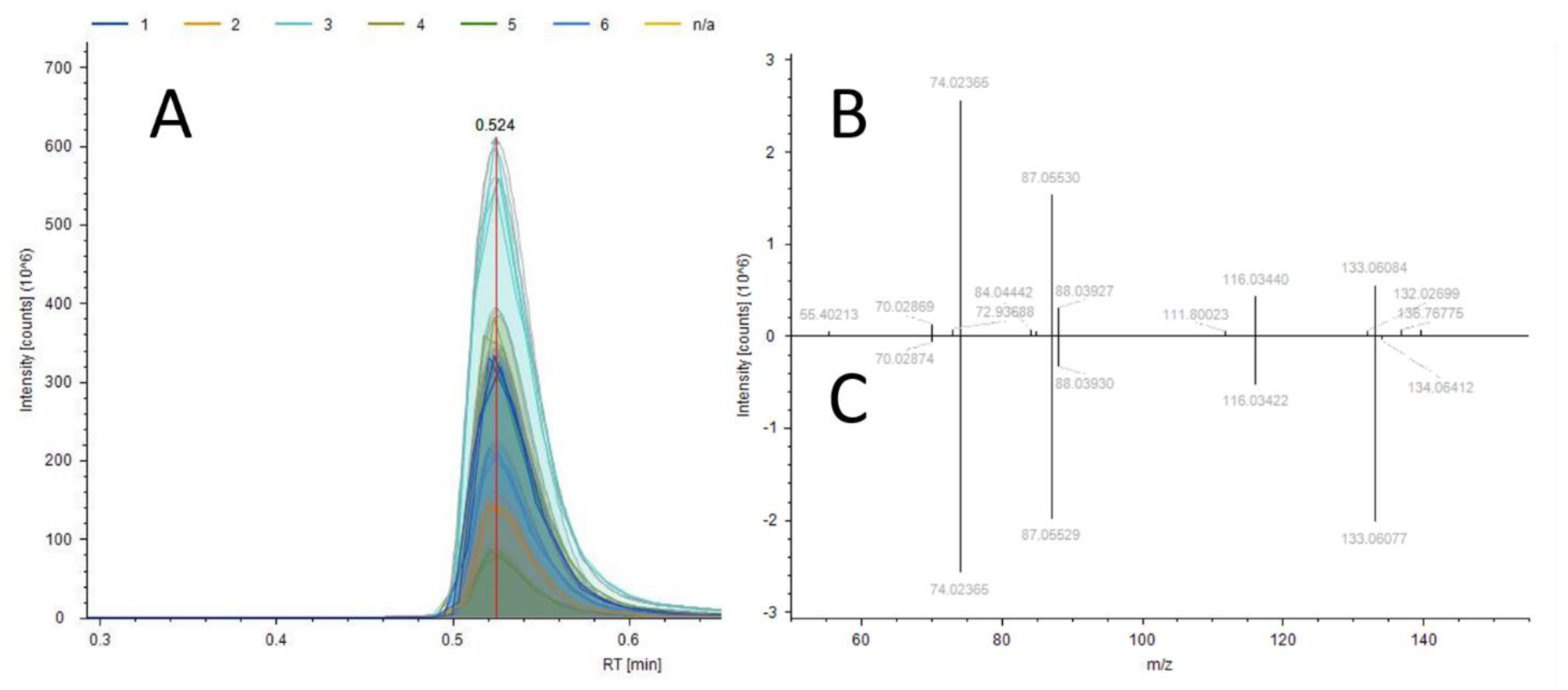
**A)** A representative extracted chromatogram for the amino acid asparagine demonstrating the peak extracted from every positive sample, with “n/a” denoting the water blank samples used for background subtraction. B) The experimental MS/MS spectra used for compound identification. C) The library MS/MS spectra from the mzCloud library for asparagine, demonstrating every fragment ion within a 3ppm mass accuracy window

**Figure 2.**
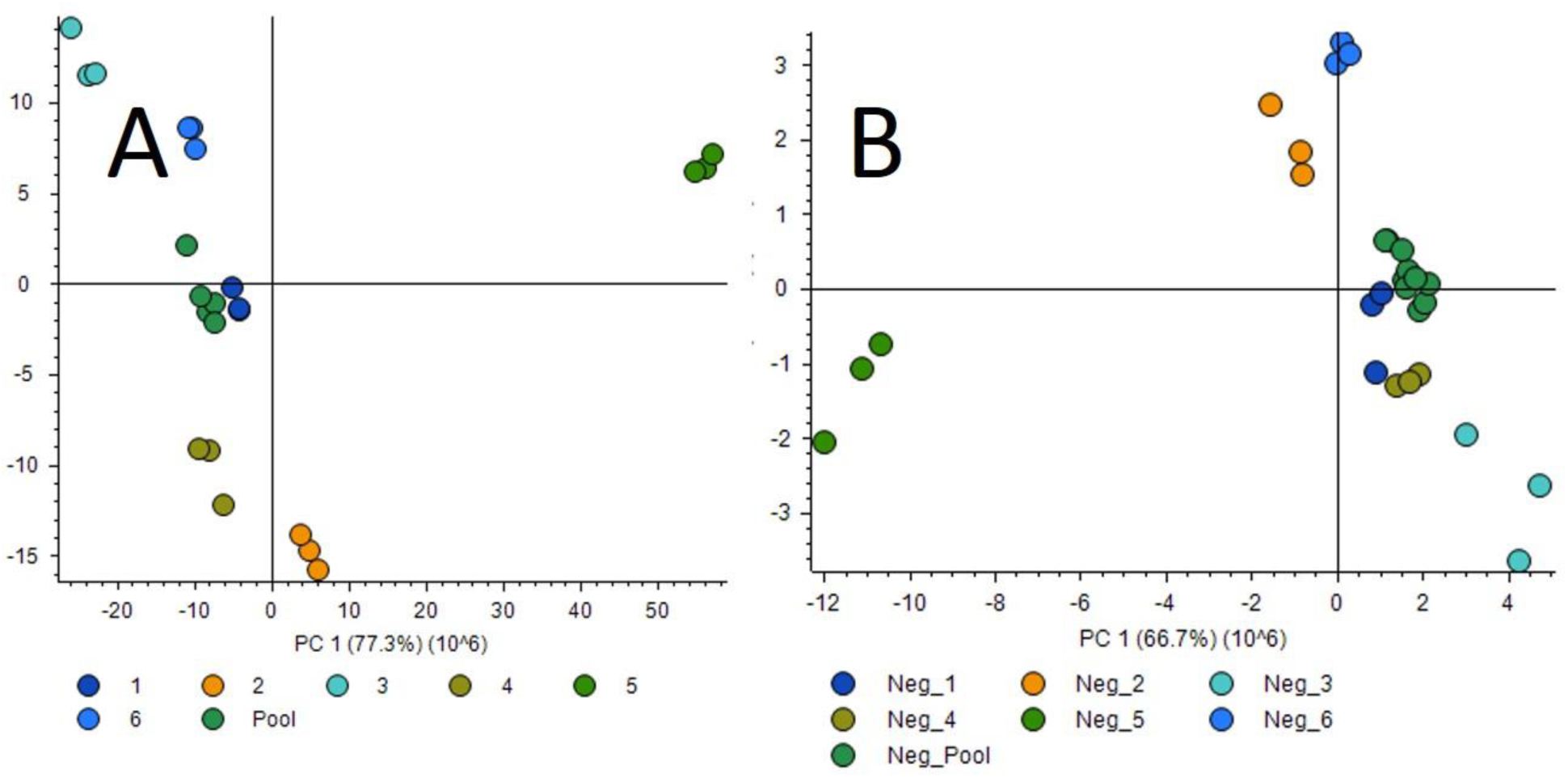
**A)** PCA Analysis of the known samples in positive. **B)** The same for the samples ran with negative polarity.

**Figure 3.**
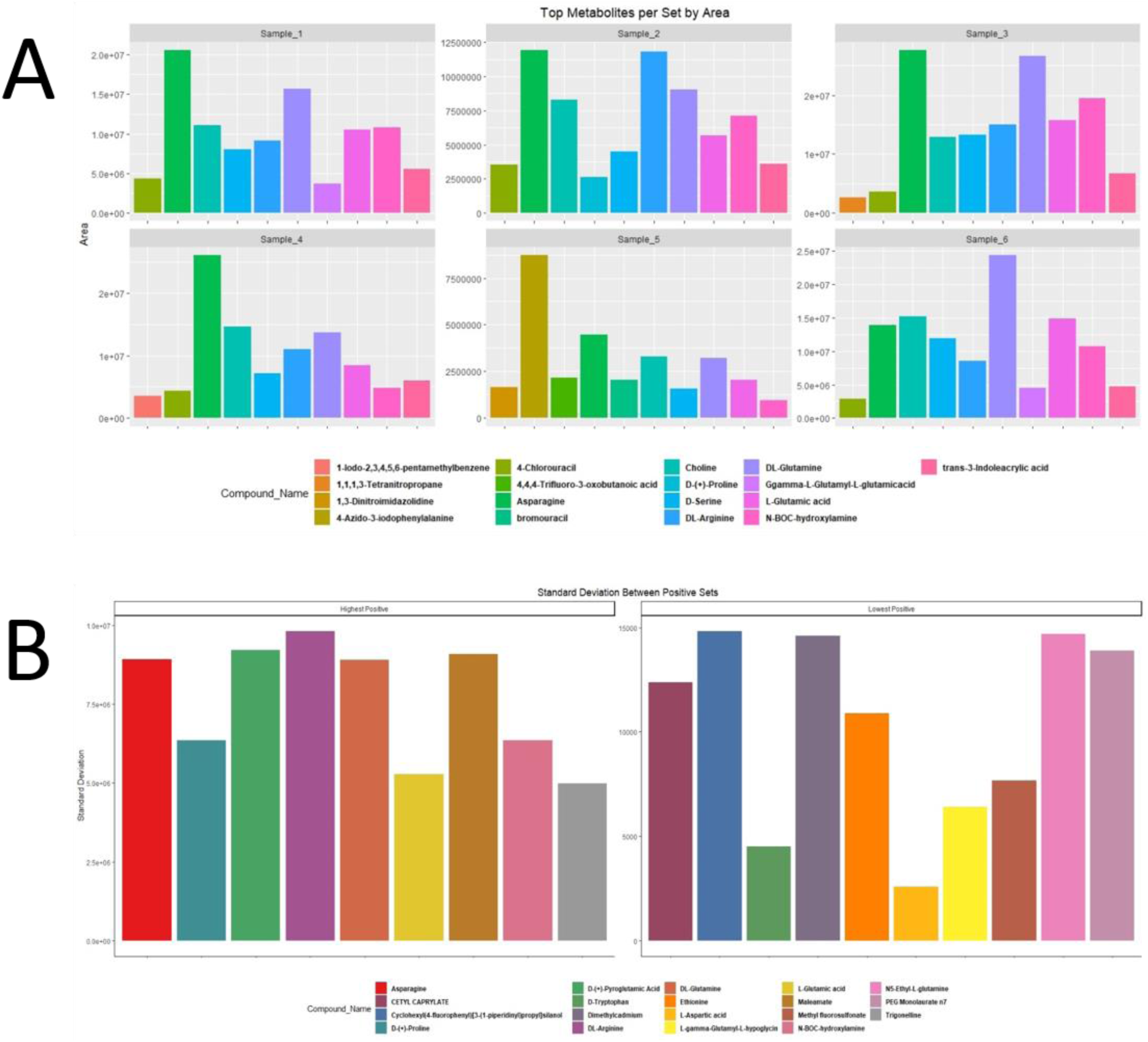
**A)** Most abundant compound identified by area using the established libraries for each sample. **B)** The compounds that exhibit the highest(left) and lowest (right) standard deviation amongst all positive files.

### Metabolite distribution

We utilized the group areas from the named compounds from each experiment polarity to determine the most abundant compounds identified for each sample. Figure 3A is a summary of the results for the positive polarity. The same analysis is repeated in the negative analysis in Supplemental Figure 1. Similar to described in previous studies, the highest abundance compounds alone and their relative quantities may be sufficient for plant material identification.^20^ In order to obtain a chemical fingerprint from each plant, two groups of small molecule compounds are initially investigated. The compounds that are conserved between each of the groups and the compounds that are indicative of unique characteristics of each sample. **Figure 3A** is a graphical representation of the metabolites that have the largest area in each of the samples. This group is made up of 14 different chemical species. Asparagine is consistently one of the highest abundant compounds found in all samples identified, also shown in **Figure 1A**, but the concentration of other chemical species can show important differences between the samples. For example, sample 4 has a chemical signature of 4-Azido-3-iodophenylalanine that is not present in any of the other samples and sample 6 and appears to be one of the most abundant species when compared to the other nine most abundant compounds. It is worth noting, however, that the factors such as ionization efficiency and relative resistance to in-source decay confounds direct measurements of abundance between compounds.^21^ Another example of a potential identifying compound is shown in Sample 3 which demonstrates 1,1,1,3-Tetranitropropane in its top ten features, something that no other set contains. Increasing the number and confidence of compound identifications with follow-up analyses will only increase the power of this approach.

### Variation of Chemical Composition Between Samples

Another important aspect of identification is observation of what each of the samples holds constant. A variation analysis was performed on the data obtained from the samples, showing the compounds that show the greatest and least variation between the samples as shown in **Figure 3B.**

### Principal Component Analysis (PCA)

The reduction of multiple variables to their principal components in reduced dimensions is a powerful and commonly utilized mechanism in many fields of science. While the main use of PCA is the identification of outliers in biological and technical replicate data, that quality of the clustering demonstrated in **Figure 2A** and **2B** suggested that we could use also use this tool as an identification method for unknown samples. To test this, identical samples were prepared separately, with the second set double blinded by an assistant. To replicate more accurate unknown sample conditions, the LC-MS instrument was operated for two days by other researchers on unrelated projects prior to running the unknown samples. As shown in **Figure 4B**, the new samples clustered with known technical replicates acquired and, in all 4 cases, with their correct sample groups. An unknown sample that was prepped and not included in the library generation analysis analysis did not cluster with any group. This sample appears to support the validity of our approach, although a higher number of samples will be required to fully validate these assays. These results appear to indicate that 11-minute UHPLC-LC-MS runs could be used to identify strains and to detect counterfeit or other adulterated samples. The main requirement would be that the strains had been previously profiled to create the library for downstream sample matching. Data reduction for the libraries themselves is currently being explored, as the experiment shown in Figure 4B requires nearly 90 minutes to run on a custom high performance server and requires nearly 40GB of storage space for each processing iteration.

**Figure 4.**
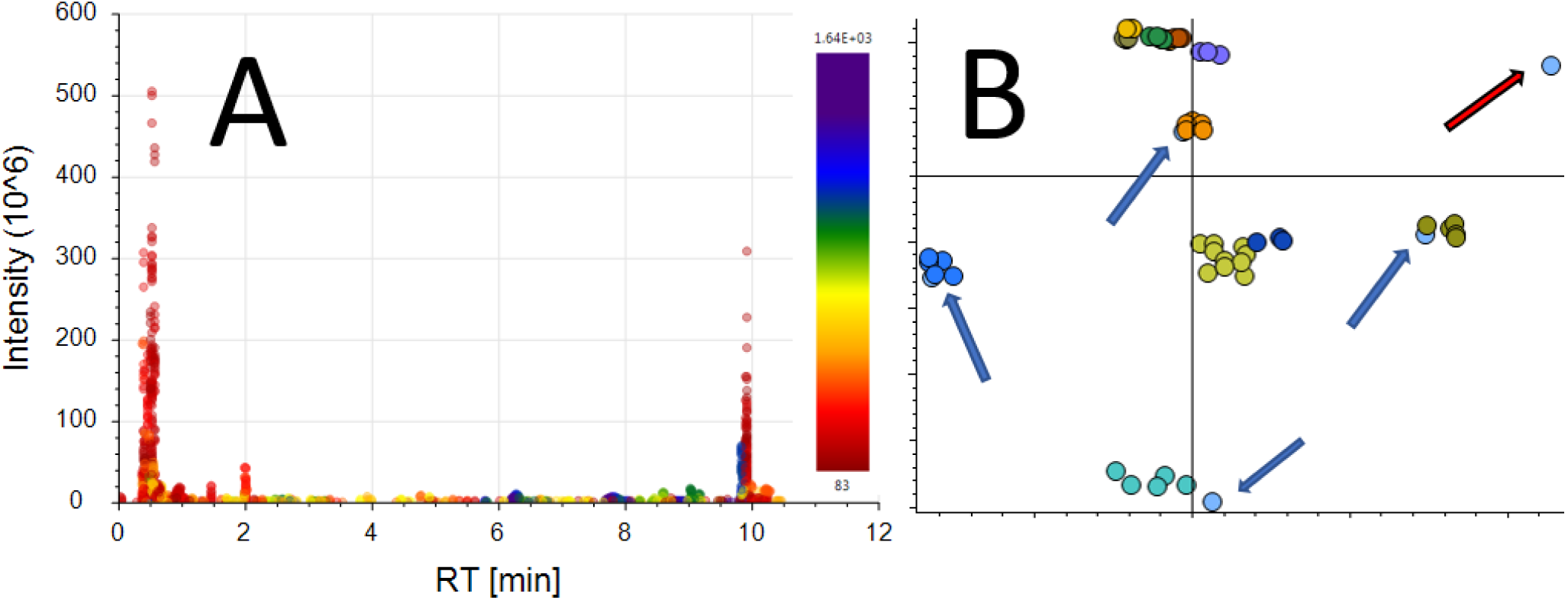
**A)** Scatterplot of a single representative RAW file. Each compound identified in the file is represented by a single point. The X axis is the compound elution time. Y is the intensity of each compound identified. The color represents the molecular weight of each compound where increasing color frequency corresponds to molecular weight. **B)** PCA plot with known and unknown samples. Blue arrows indicate unknown samples that cluster with their library known counterparts. The red arrow indicates a sample that was not included in this analysis and represents a counterfeit extract

**Figure 5.**
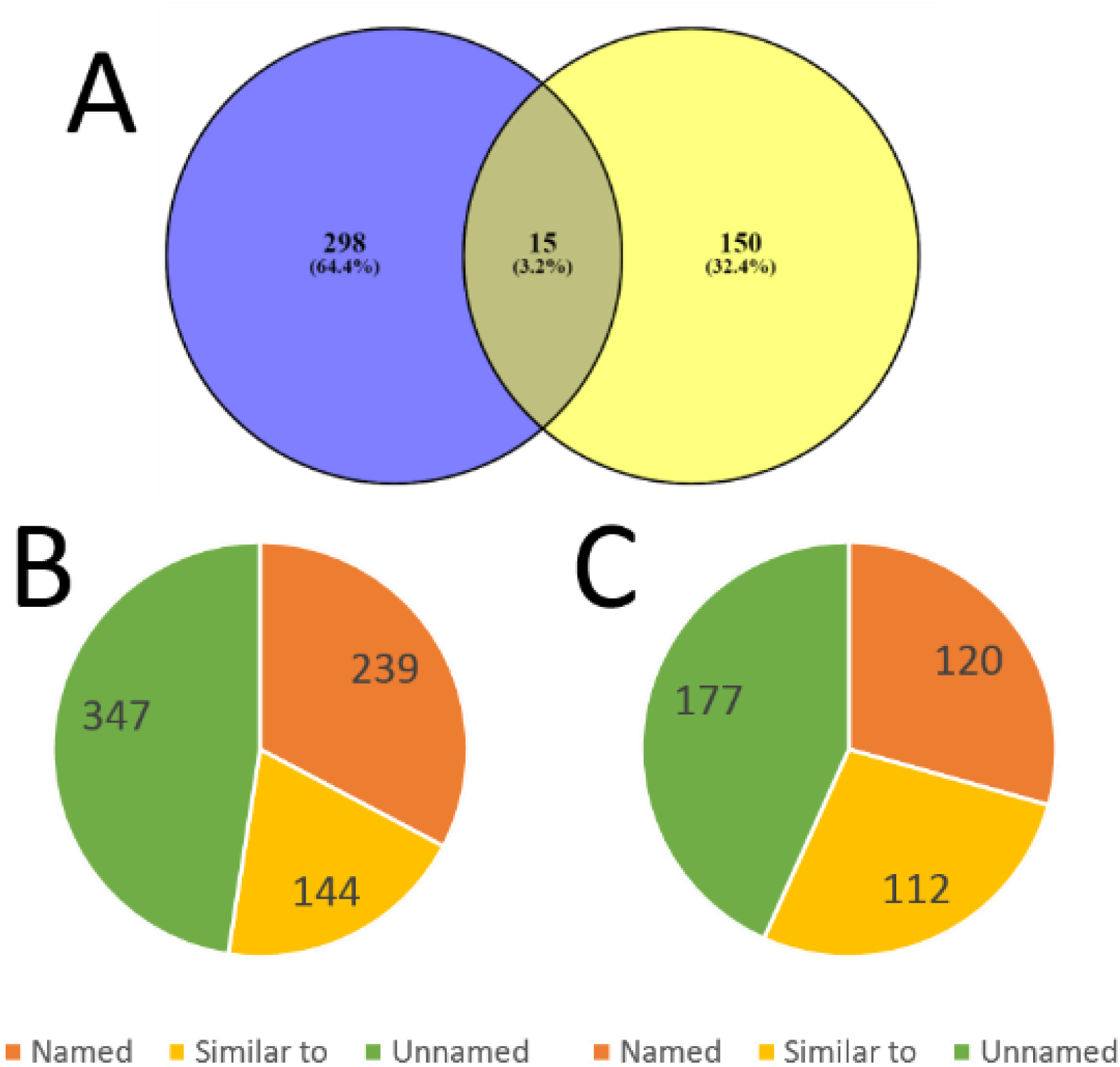
**A)** The overlap of named compounds identified in positive mode (left) and negative mode (right). **B)** The distribution of feature identifications in positive mode **C)** Same as B for negative ionization mode

## Conclusions

We have utilized tools developed for global metabolomic profiling to develop high resolution chemical profiles for multiple medical Cannabis products. We find that the majority of small molecule features identified cannot be assigned chemical identifications from commonly utilized small molecule libraries. However, ultra-high resolution mass alone was sufficient to generate putative molecular formulas for nearly 90% of all identified features. Using the generated chemical profiles, reliant on retention time and mass alone, we find that relatively simple statistical tools can be rapidly used to identify unknown samples. We conclude that the use of rapid UHPLC high resolution mass spectrometry assays will be a valuable tool for the identification of counterfeit and adulterated medical products of even the highest complexity. The materials utilized in this study were chosen due as an example of the highest complexity medical products in use today. The surprising number of unknown compounds present in these medical materials will be the focus of follow-up work to be detailed elsewhere.

## References

(1) Lommen, A.; Gerssen, A.; Oosterink, J. E.; Kools, H. J.; Ruiz-Aracama, A.; Peters, R. J. B.; Mol, H. G. J. Ultra-Fast Searching Assists in Evaluating Sub-Ppm Mass Accuracy Enhancement in UHPLC/Orbitrap MS Data. Metabolomics 2011. https://doi.org/10.1007/s11306-010-0230-y.

(2) Martin, J. C.; Maillot, M.; Mazerolles, G.; Verdu, A.; Lyan, B.; Migné, C.; Defoort, C.; Canlet, C.; Junot, C.; Guillou, C.; et al. Can We Trust Untargeted Metabolomics? Results of the Metabo-Ring Initiative, a Large-Scale, Multi-Instrument Inter-Laboratory Study. Metabolomics 2015. https://doi.org/10.1007/s11306-014-0740-0.

(3) Yang, X.; Neta, P.; Stein, S. E. Quality Control for Building Libraries from Electrospray Ionization Tandem Mass Spectra. Anal. Chem. 2014. https://doi.org/10.1021/ac500711m.

(4) Kind, T.; Tsugawa, H.; Cajka, T.; Ma, Y.; Lai, Z.; Mehta, S. S.; Wohlgemuth, G.; Barupal, D. K.; Showalter, M. R.; Arita, M.; et al. Identification of Small Molecules Using Accurate Mass MS/MS Search. Mass Spectrometry Reviews. 2018. https://doi.org/10.1002/mas.21535.

(5) Johnson, R. Food Fraud and “Economically Motivated Adulteration” of Food and Food Ingredients; 2014. https://doi.org/10.1177/014860717700100306.

(6) Spink, J.; Moyer, D. C. Defining the Public Health Threat of Food Fraud. J. Food Sci. 2011. https://doi.org/10.1111/j.1750-3841.2011.02417.x.

(7) van Ruth, S. M.; Huisman, W.; Luning, P. A. Food Fraud Vulnerability and Its Key Factors. Trends in Food Science and Technology. 2017. https://doi.org/10.1016/j.tifs.2017.06.017.

(8) Elder, D. The Cost of Drug Counterfeiting. European Pharmaceutical Review. 2015.

(9) Pichini, S.; Solimini, R.; Berretta, P.; Pacifici, R.; Busardò, F. P. Acute Intoxications and Fatalities From Illicit Fentanyl and Analogues: An Update. Therapeutic drug monitoring. 2018. https://doi.org/10.1097/FTD.0000000000000465.

(10) Samms, W. C.; Jiang, Y. J.; Dixon, M. D.; Houck, S. S.; Mozayani, A. Analysis of Alprazolam by DART-TOF Mass Spectrometry in Counterfeit and Routine Drug Identification Cases. J. Forensic Sci. 2011. https://doi.org/10.1111/j.1556-4029.2011.01767.x.

(11) Arens, A. M.; Van Wijk, X. M. R.; Vo, K. T.; Lynch, K. L.; Wu, A. H. B.; Smollin, C. G. Adverse Effects from Counterfeit Alprazolam Tablets. JAMA Internal Medicine. 2016. https://doi.org/10.1001/jamainternmed.2016.4306.

(12) Mead, A. The Legal Status of Cannabis (Marijuana) and Cannabidiol (CBD) under U.S. Law. Epilepsy and Behavior. 2017. https://doi.org/10.1016/j.yebeh.2016.11.021.

(13) Grucza, R. A.; Vuolo, M.; Krauss, M. J.; Plunk, A. D.; Agrawal, A.; Chaloupka, F. J.; Bierut, L. J. Cannabis Decriminalization: A Study of Recent Policy Change in Five U.S. States. Int. J. Drug Policy 2018. https://doi.org/10.1016/j.drugpo.2018.06.016.

(14) Oliveros, J. C. Venny. An interactive tool for comparing lists with Venn Diagrams.

(15) Burke, M. C.; Mirokhin, Y. A.; Tchekhovskoi, D. V.; Markey, S. P.; Heidbrink Thompson, J.; Larkin, C.; Stein, S. E. The Hybrid Search: A Mass Spectral Library Search Method for Discovery of Modifications in Proteomics. J. Proteome Res. 2017. https://doi.org/10.1021/acs.jproteome.6b00988.

(16) Kind, T.; Fiehn, O. Seven Golden Rules for Heuristic Filtering of Molecular Formulas Obtained by Accurate Mass Spectrometry. BMC Bioinformatics 2007. https://doi.org/10.1186/1471-2105-8-105.

(17) Broadhurst, D. I.; Kell, D. B. Statistical Strategies for Avoiding False Discoveries in Metabolomics and Related Experiments. Metabolomics 2006. https://doi.org/10.1007/s11306-006-0037-z.

(18) Wang, X.; Jones, D. R.; Shaw, T. I.; Cho, J. H.; Wang, Y.; Tan, H.; Xie, B.; Zhou, S.; Li, Y.; Peng, J. Target-Decoy-Based False Discovery Rate Estimation for Large-Scale Metabolite Identification. J. Proteome Res. 2018. https://doi.org/10.1021/acs.jproteome.8b00019.

(19) Scheubert, K.; Hufsky, F.; Petras, D.; Wang, M.; Nothias, L. F.; Dührkop, K.; Bandeira, N.; Dorrestein, P. C.; Böcker, S. Significance Estimation for Large Scale Metabolomics Annotations by Spectral Matching. Nat. Commun. 2017. https://doi.org/10.1038/s41467-017-01318-5.

(20) Berman, P.; Futoran, K.; Lewitus, G. M.; Mukha, D.; Benami, M.; Shlomi, T.; Meiri, D. A New ESI-LC/MS Approach for Comprehensive Metabolic Profiling of Phytocannabinoids in Cannabis. Sci. Rep. 2018. https://doi.org/10.1038/s41598-018-32651-4.

(21) CS Ho, CWK Lam*, MHM Chan, RCK Cheung, LK Law, LCW Lit, KF Ng, MWM Suen, and H. T. Electrospray Ionisation Mass Spectrometry: Principles and Clinical Applications. Clin Biochem Rev 2003. https://doi.org/10.1146/annurev.bi.64.070195.001531.

