## Supplemental Table 2 for "Application of Global Metabolomics to the Identification of Complex Counterfeit Medicinal Products"

Search name: 2_19_19_Flower_Untargeted_Positive-first_runs

Search description: Untargeted food research workflow with statistics: Find and identify unknowns including the differences between sample groups.

- Performs retention time alignment, unknown compound detection, and compound grouping across all samples. Predicts elemental compositions for all compounds, fills gaps across all samples, and hides chemical background (using Blank samples). Identifies compounds using mzCloud (ddMS2), ChemSpider (exact mass or formula) and local database searches against Mass Lists (exact mass and RT) and mzVault spectral libraries. Performs similarity search for all compounds with ddMS2 data using mzCloud. Calculates differential analysis (t-test or ANOVA), determines p-values, adjusted p-values, ratios, fold change, CV, etc.).

Search date: 2/19/2019 9:08:46 AM

Created with Discoverer version: 2.1.0.398

[Input Files (0)]

-->Select Spectra (1)

[Select Spectra (1)]

-->Align Retention Times (35)

[Align Retention Times (35)]

-->Detect Unknown Compounds (25)

[Detect Unknown Compounds (25)]

-->Group Unknown Compounds (24)

-->Merge Features (14)

[Group Unknown Compounds (24)]

-->Search mzCloud (27)

-->Fill Gaps (38)

-->Predict Compositions (26)

-->Search ChemSpider (22)

-->Search Mass Lists (23)

[Fill Gaps (38)]

-->Mark Background Compounds (37)

[Predict Compositions (26)]

-->Search ChemSpider (22)

[Search mzCloud (27)]

[Mark Background Compounds (37)]

[Search ChemSpider (22)]

[Search Mass Lists (23)]

[Merge Features (14)]

[Assign Compound Annotations (29)]

[Differential Analysis (30)]

------------------------------------------------------------------

Processing node 0: Input Files

------------------------------------------------------------------

Input Data:

- File Name(s) (Hidden):

------------------------------------------------------------------

Processing node 1: Select Spectra

------------------------------------------------------------------

1. General Settings:

- Precursor Selection: Use MS(n - 1) Precursor

- Use New Precursor Reevaluation: True

- Use Isotope Pattern in Precursor Reevaluation: True

- Store Chromatograms: False

2. Spectrum Properties Filter:

- Lower RT Limit: 0

- Upper RT Limit: 0

- First Scan: 0

- Last Scan: 0

- Ignore Specified Scans: (not specified)

- Lowest Charge State: 0

- Highest Charge State: 0

- Min. Precursor Mass: 100 Da

- Max. Precursor Mass: 5000 Da

- Total Intensity Threshold: 0

- Minimum Peak Count: 1

3. Scan Event Filters:

- Mass Analyzer: (not specified)

- MS Order: Any

- Activation Type: (not specified)

- Min. Collision Energy: 0

- Max. Collision Energy: 1000

- Scan Type: Any

- Polarity Mode: (not specified)

4. Peak Filters:

- S/N Threshold (FT-only): 1.5

5. Replacements for Unrecognized Properties:

- Unrecognized Charge Replacements: 1

- Unrecognized Mass Analyzer Replacements: ITMS

- Unrecognized MS Order Replacements: MS2

- Unrecognized Activation Type Replacements: CID

- Unrecognized Polarity Replacements: +

- Unrecognized MS Resolution@200 Replacements: 60000

- Unrecognized MSn Resolution@200 Replacements: 30000

------------------------------------------------------------------

Processing node 35: Align Retention Times

------------------------------------------------------------------

1. General Settings:

- Alignment Model: Adaptive curve

- Alignment Fallback: Use Linear Model

- Maximum Shift [min]: 2

- Shift Reference File: True

- Mass Tolerance: 5 ppm

- Remove Outlier: True

------------------------------------------------------------------

Processing node 25: Detect Unknown Compounds

------------------------------------------------------------------

1. General Settings:

- Mass Tolerance [ppm]: 5 ppm

- Intensity Tolerance [%]: 30

- S/N Threshold: 3

- Min. Peak Intensity: 1000000

- Ions:

[2M+ACN+H]+1

[2M+ACN+Na]+1

[2M+FA-H]-1

[2M+H]+1

[2M+K]+1

[2M+Na]+1

[2M+NH4]+1

[2M-H]-1

[2M-H+HAc]-1

[M+2H]+2

[M+3H]+3

[M+ACN+2H]+2

[M+ACN+H]+1

[M+ACN+Na]+1

[M+Cl]-1

[M+DMSO+H]+1

[M+FA-H]-1

[M+H]+1

[M+H+K]+2

[M+H+MeOH]+1

[M+H+Na]+2

[M+H+NH4]+2

[M+H-H2O]+1

[M+H-NH3]+1

[M+K]+1

[M+Na]+1

[M+NH4]+1

[M-2H]-2

[M-2H+K]-1

[M-H]-1

[M-H+HAc]-1

[M-H+TFA]-1

[M-H-H2O]-1

- Base Ions: [M+H]+1; [M-H]-1

- Min. Element Counts: C H

- Max. Element Counts: C90 H190 Br3 Cl4 F6 K2 N10 Na2 O18 P3 S5

2. Peak Detection:

- Filter Peaks: True

- Max. Peak Width [min]: 0.8

- Remove Singlets: False

- Min. # Scans per Peak: 3

- Min. # Isotopes: 1

------------------------------------------------------------------

Processing node 24: Group Unknown Compounds

------------------------------------------------------------------

1. Compound Consolidation:

- Mass Tolerance: 5 ppm

- RT Tolerance [min]: 0.1

2. Fragment Data Selection:

- Preferred Ions: [M+H]+1; [M-H]-1

------------------------------------------------------------------

Processing node 27: Search mzCloud

------------------------------------------------------------------

1. Search Settings:

- Compound Classes: All

- Match Ion Activation Type: True

- Match Ion Activation Energy: Match with Tolerance

- Ion Activation Energy Tolerance: 20

- Apply Intensity Threshold: True

- Precursor Mass Tolerance: 10 ppm

- FT Fragment Mass Tolerance: 10 ppm

- IT Fragment Mass Tolerance: 0.4 Da

- Identity Search: Cosine

- Similarity Search: Similarity Forward

- Library: Reference

- Post Processing: Recalibrated

- Match Factor Threshold: 50

- Max. # Results: 20

------------------------------------------------------------------

Processing node 38: Fill Gaps

------------------------------------------------------------------

1. General Settings:

- Mass Tolerance: 5 ppm

- S/N Threshold: 1.5

- Use Real Peak Detection: True

------------------------------------------------------------------

Processing node 37: Mark Background Compounds

------------------------------------------------------------------

1. General Settings:

- Max. Sample/Blank: 5

- Max. Blank/Sample: 0

- Hide Background: True

------------------------------------------------------------------

Processing node 26: Predict Compositions

------------------------------------------------------------------

1. Prediction Settings:

- Mass Tolerance: 5 ppm

- Min. Element Counts: C H

- Max. Element Counts: C90 H190 Br3 Cl4 F6 N10 O18 P3 S5

- Min. RDBE: 0

- Max. RDBE: 40

- Min. H/C: 0.1

- Max. H/C: 3.5

- Max. # Candidates: 10

- Max. # Internal Candidates: 200

2. Pattern Matching:

- Intensity Tolerance [%]: 30

- Intensity Threshold [%]: 0.1

- S/N Threshold: 3

- Min. Spectral Fit [%]: 30

- Min. Pattern Cov. [%]: 90

- Use Dynamic Recalibration: True

3. Fragments Matching:

- Use Fragments Matching: True

- Mass Tolerance: 5 ppm

- S/N Threshold: 3

------------------------------------------------------------------

Processing node 22: Search ChemSpider

------------------------------------------------------------------

1. Search Settings:

- Mass Tolerance: 5 ppm

- Database(s):

ACToR: Aggregated Computational Toxicology Resource

FDA UNII - NLM

FooDB

- Max. # of results per compound: 100

- Max. # of Predicted Compositions to be searched per Compound: 3

- Result Order (for Max. # of results per compound): Order By Reference Count (DESC)

2. Predicted Composition Annotation:

- Check All Predicted Compositions: True

------------------------------------------------------------------

Processing node 23: Search Mass Lists

------------------------------------------------------------------

1. Search Settings:

- Input file(s): EFS HRAM Compound Database.csv

- Mass Tolerance: 5 ppm

- Show extra Fields as Columns: False

- Consider Retention Time: True

- RT Tolerance : 0.05

------------------------------------------------------------------

Processing node 14: Merge Features

------------------------------------------------------------------

1. Peak Consolidation:

- Mass Tolerance: 5 ppm

- RT Tolerance [min]: 0.1

------------------------------------------------------------------

Processing node 29: Assign Compound Annotations

------------------------------------------------------------------

1. General Settings:

- Mass Tolerance: 5 ppm

2. Data Sources:

- Data Source #1: mzCloud Search

- Data Source #2: Predicted Compositions

- Data Source #3: MassList Match

- Data Source #4: ChemSpider Search

------------------------------------------------------------------

Processing node 30: Differential Analysis

------------------------------------------------------------------

1. General Settings:

- Log10 Transform Values: True
