## Supplementary figures and images for "Application of Global Metabolomics to the Identification of Complex Counterfeit Medicinal Products"

### Supplemental Figure 1

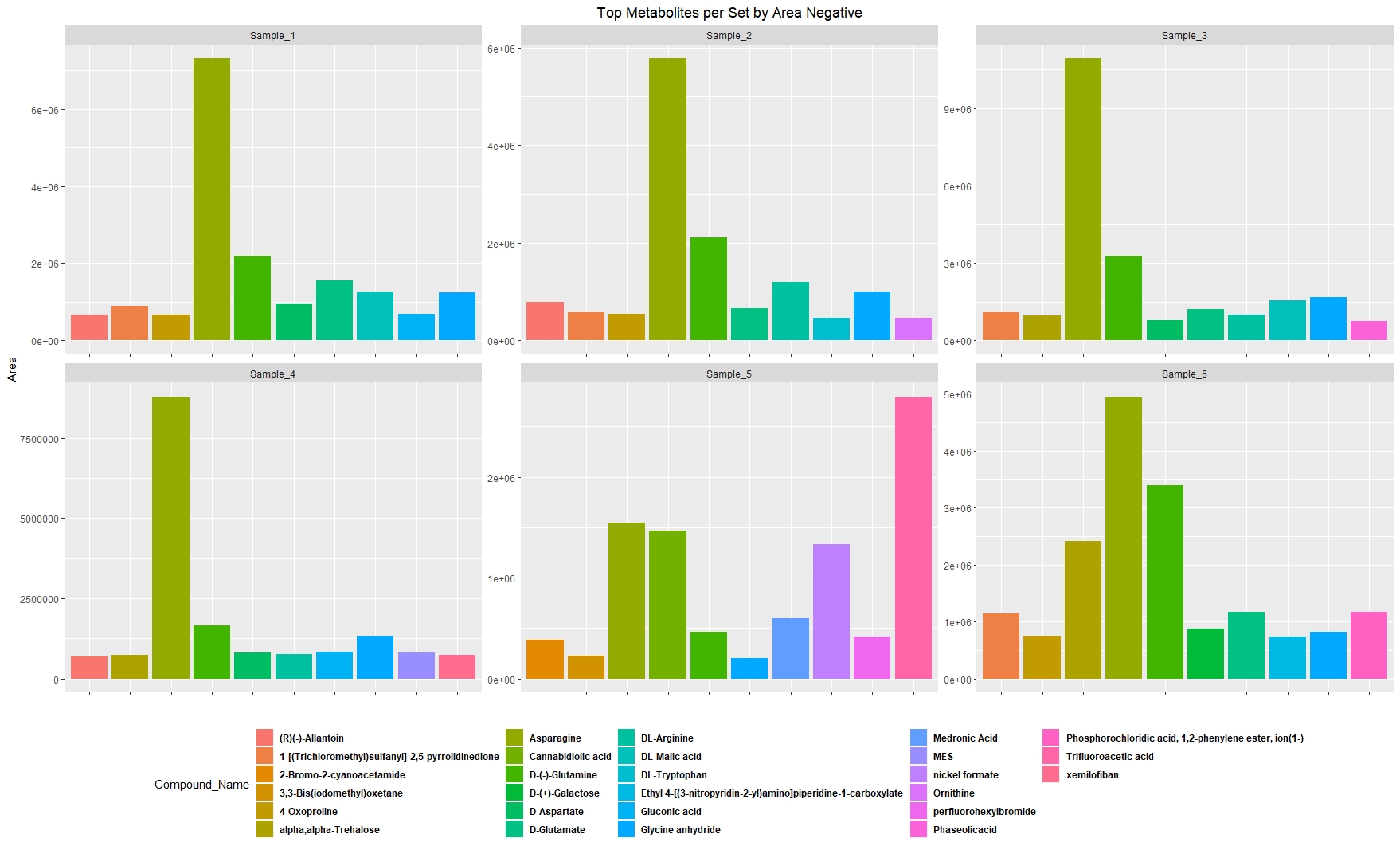
